# *De novo* transcriptome assembly and genome annotation of the fat-tailed dunnart (*Sminthopsis crassicaudata*)

**DOI:** 10.1101/2023.11.16.567318

**Authors:** Neke Ibeh, Charles Y. Feigin, Stephen R. Frankenberg, Davis J. McCarthy, Andrew J. Pask, Irene Gallego Romero

## Abstract

Marsupials exhibit highly specialized patterns of reproduction and development, making them uniquely valuable for comparative genomics studies with their sister lineage, eutherian (also known as placental) mammals. However, marsupial genomic resources still lag far behind those of eutherian mammals, limiting our insight into mammalian diversity. Here, we present a series of novel genomic resources for the fat-tailed dunnart (*Sminthopsis crassicaudata*), a mouse-like marsupial that, due to its ease of husbandry and *ex-utero* development, is emerging as a laboratory model. To enable wider use, we have generated a multi-tissue *de novo* transcriptome assembly of dunnart RNA-seq reads spanning 12 tissues. This highly representative transcriptome is comprised of 2,093,982 assembled transcripts, with a mean transcript length of 830 bp. The transcriptome mammalian BUSCO completeness score of 93% is the highest amongst all other published marsupial transcriptomes. Additionally, we report an improved fat-tailed dunnart genome assembly which is 3.23 Gb long, organized into 1,848 scaffolds, with a scaffold N50 of 72.64 Mb. The genome annotation, supported by assembled transcripts and *ab initio* predictions, revealed 21,622 protein-coding genes. Altogether, these resources will contribute greatly towards characterizing marsupial biology and mammalian genome evolution.

## Data Description

### Background and context

Marsupials are a strikingly diverse mammalian group predominantly found in Australasia (Australia, Tasmania, New Guinea, and nearby islands), with a number of species also present in the Americas [4, 5]. While many marsupials are convergent in form to eutherian mammals [6, 7, 8, 9, 10, 11, 12, 13, 14], their adaptations to the niches they occupy include vastly specialized physiology [15, 16, 17, 18, 19, 20, 21], behaviour [22, 23, 24], and modes of reproduction [25, 26, 27, 28, 29, 30], thereby offering a unique perspective on mammalian diversity and life-cycles. To date, marsupial studies have significantly contributed towards elucidating various aspects of mammalian biology, including: reproductive physiology [27, 28, 29, 30], sex determination [31, 32, 33, 34, 35, 36, 37], X-chromosome inactivation [38, 39, 40, 41, 27], age-related obesity [42], postnatal development [43, 44, 18, 19, 20, 21, 45], and genome evolution [46, 47, 48, 49, 50, 51, 52, 53], to name a few. Consequently, marsupials represent a critical comparative model system through which mammalian biology can be better understood.

In spite of the importance of well-developed marsupial models, marsupial genomic resources still lag far behind those of their eutherian counterparts. Currently, only seven marsupial species have an annotated genome available: the gray short-tailed opossum (*Monodelphis domestica*) [54], the tammar wallaby (*Macropus eugenii*) [55], the Tasmanian devil (*Sarcophilus harrisii*) [46], the brown antechinus (*Antechinus stuartii*) [56], the koala (*Phascolarctos cinereus*) [57], the numbat (*Myrmecobius fasciatus*) [58], and the eastern quoll (*Dasyurus viverrinus*) [59], with genome assembly recovery of complete single-copy mammalian BUSCOs (Benchmarking Universal Single-Copy Orthologs) ranging from 73.1% to 92.4% [59]. The global transcriptomes generated for some of these species have BUSCO scores ranging from 76.4% to 84% [56, 58]. Annotated genomes and global transcriptomes are of paramount importance for attaching biological meaning to sequencing data.

Recently, the fat-tailed dunnart (*Sminthopsis crassicaudata*) has emerged as a key laboratory marsupial model for understanding mammalian development and evolution [60, 61, 62, 63, 64, 45]. A nocturnal species belonging to the family Dasyuridae, the fat-tailed dunnart has adapted to a wide range of habitats and can be found across south and central mainland Australia [65] (Figure 1A and B). As one of the smallest carnivorous marsupials, adults weigh an average of 15 grams. Fat-tailed dunnarts exhibit some of the shortest known gestation times for mammals (13 days), with much of their development occurring postnatally. Fat-tailed dunnart neonates reside in their mother’s pouch, thereby allowing continuous and non-invasive experimental access [66, 67]. The extremely altricial state of the dunnart young, along with very simple husbandry requirements, have facilitated the dunnart’s role as a model species for comparative mammalian studies and conservation strategies. Nonetheless, the paucity of genomic resources for the fat-tailed dunnart has limited our understanding of this species at the gene level.

**Figure 1.**
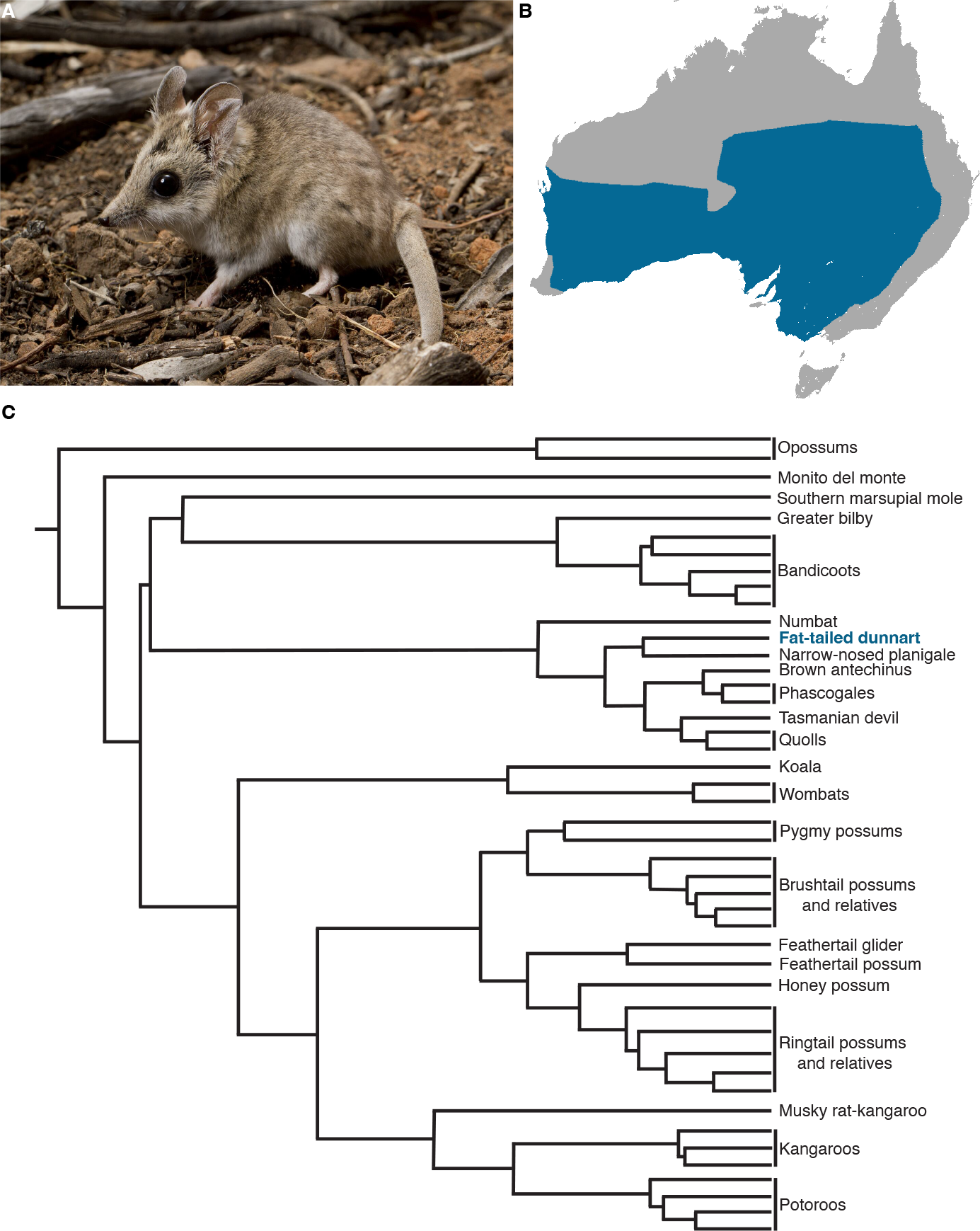
The fat-tailed dunnart (*Sminthopsis crassicaudata*). **(A)** Adult fat-tailed dunnart captured in Ned’s Corner, Victoria (Photo credit: David Paul, Museums Victoria). **(B)** The fat-tailed dunnart’s range across Australia (CC BY-SA 3.0)[1]. **(C)** Phylogeny of extant marsupial orders (based on [2] and [3]). The fat-tailed dunnart (blue font) is a member of the order Dasyuromorphia.

Here, to address this knowledge gap, we present a comprehensive transcriptome and annotation for the fat-tailed dunnart, supplementing an improved draft genome assembly. This annotation effort, made possible through a multi-tissue transcriptome assembly, yielded 21,622 protein-coding genes. The global transcriptome had a 93.3% recovery of complete mammalian BUSCOs (Benchmarking Universal Single-Copy Orthologs). This first-draft annotation and global transcriptome can serve as tools with which the genomic architecture of the fat-tailed dunnart, an emerging marsupial model species, can be better understood. Most importantly, these comprehensive resources contribute to the growing body of research on marsupial genomics and are therefore invaluable tools for future mammalian studies.

## Methods

### Draft genome assembly

Fat-tailed dunnart ONT (∽171 Gb) and Pacific Biosciences CRL (∽18 Gb) long reads [68] and Illumina short reads (∽447.5 Gb in 2×150 bp format) [53] were combined to produce an improved draft genome assembly (v1.1). Briefly, *de novo* contigs were first assembled from long reads *≥* 10 kilobases using Flye v2.9 [69] (parameters: –pacbio-raw, –genome-size 3g –iterations 2 –scaffold). Uncollapsed haplotypes were removed using purge_dups [70] with automatic coverage threshold detection. A second round of scaffolding was then performed using LongStitch v1.0.1 [71] (mode: ntLink-arks with estimated genome size of 3 Gb). The resulting assembly was then polished in two rounds using Pilon v1.24 [72] (parameters: –vcf –diploid –chunksize 10000000 –fix snps,indels,gaps –minqual 15). To do this, Illumina short reads were first filtered and trimmed with Trimmomatic v0.38 [73] (parameters: SLIDINGWINDOW:5:30, MINLEN:75, AVGQUAL:30). Reads were then aligned against the assembly using BWA-MEM2 [74] (parameter: –M) and the resulting alignments were filtered with samtools view v1.11 [75] (parameters: –h –b –q 30 –F 3340 –f 3). Benchmark mammal ortholog recovery for the assembly was determined using BUSCO v5.2.2, in genome mode, using the Mammalia_odb v10 database of orthologs (9226 BUSCOs).

### Sample collection and sequencing

Adult and fetal fat-tailed dunnart tissues were collected from several individuals for short-read Illumina sequencing. Tissues included: late pregnancy allantois (n=3), amnion (n=3), distal yolk sac without vasculature (n=2), proximal yolk sac with vasculature (n=2), endometrium (n=4), ovary (n=3), oviduct (n=2), combined uterus and oviduct (n=1), testis (n=1), liver (n=1), eye (n=1), and prostate gland (n=1).

Multiple RNA samples were pooled in approximately equal proportions for Iso-Seq, namely, allantois, amnion, distal and proximal yolk sacs, endometrium, oviduct, endometrium, ovary, testis, liver, eye, gastrula-stage conceptus, and late fetus. All RNA samples were extracted using Qiagen RNeasy Mini or Micro kits according to manufacturer’s instructions, with Illumina and Iso-Seq library construction and sequencing outsourced to Azenta Life Sciences (USA).

### *De novo* transcriptome assembly

We used a total of 24 dunnart RNA-seq samples, originating from various tissues (liver, testis, prostate, ovary, oviduct, uterus, eye, whole neonate, allantois, amnion, distal yolk sac, proximal yolk sac, and endometrium). Of these samples, 23 were short-read RNA-seq, with read lengths ranging from 100-150 bp, and 1 was a long-read Iso-seq sample (mean length of 5,400 bp). All samples were quality checked using FastQC v0.11.9 [76]. Quality trimming of the RNA-seq reads was carried out using Trimmomatic v0.38 [73] (parameters: SLIDINGWINDOW:4:28, MINLEN:25, AVGQUAL:28). Post trimming, 464M paired reads remained.

To generate a global dunnart transcriptome, the trimmed, paired-end RNA-seq reads were used as input to Trinity v2.13.2 [77]. We applied default *in silico* read normalization, and set the minimum assembled contig length to report to 200. Circular consensus reads were incorporated for Iso-seq long-read correction (parameter: –long_reads). Contig assembly was executed using three different k-mer settings: 25, 29, and 32. We chose these values because 25 and 32 are the minimum and maximum permitted values for the Trinity contig assembly step. Assembly statistics were obtained using the Trinity script TrinityStats.pl [77]. A reference-free evaluation of assembly quality was conducted using RSEM-EVAL, a component package of Detonate v1.11 [78]. RSEM-EVAL provides a weighted quality score using a probabilistic model. Although these scores are always negative, when comparing two assemblies, a higher value represents a higher quality assembly. The completeness of the full-length assemblies was evaluated using Benchmarking Universal Single-Copy Orthologs (BUSCO) [79]. The BUSCO gene sets are comprised of nearly universally distributed single-copy orthologous genes representing various phylogenetic levels. Here, BUSCO v5.2.2 assessment was carried out in transcriptome mode using the Mammalia_odb v10 database of orthologs.

To quantify the RNA-seq read representation of the assembly, all reads were mapped back to the global transcriptome assembly using Bowtie2 v2.4.5 [80], setting a maximum of 20 distinct alignments for each read (parameter: –k 20). Prior to annotation, transcript redundancy in the global transcriptome was reduced using CD-HIT v4.8.1 [81] with a homology threshold of 1 (parameter: –c 1) to avoid filtering out true isoforms.

### Transcriptome functional annotation

Functional annotation of the assembled transcripts was conducted using the Trinotate v3.2.2 [77] analysis protocol. First, Transdecoder [77] was used to identify all open reading frames (with a minimum length of 100 amino acids) and predict coding regions within transcripts. Sequence and domain homologies were captured by running BLAST+ v2.13.0 [82] (parameters: –max_target_seqs 1 –outfmt 6 –evalue 1e-5) against a combined protein database consisting of the UniProt/Swiss-Prot non-redundant protein sequences [83] from human (UP000005640), house mouse (UP000000589), Tasmanian devil (UP000007648), koala (UP000515140), tammar wallaby (txid9315), gray short-tailed opossum (UP000002280), and numbat (txid55782). Functional domains were identified by running a HMMer v3.3.2 [84] search against the PFAM v35.0 [85] database using the predicted protein sequences. Signal peptides and transmembrane domains were predicted using the SignalP v6.0 [86] and DeepTMHMM v1.0.24 [87] software tools, respectively.

### Genome annotation

Annotation of the dunnart draft genome was conducted using a combination of *ab initio* gene prediction algorithms and homology-based methods (Figure 2). First, genome repeats were masked using RepeatMasker v4.0.6 [88], with complex repeats being hard-masked while simple repeats were soft-masked. Preliminary gene models were constructed with MAKER2 [89] by aligning the assembled transcriptome and homologous protein sequences to the masked genome using minimap2 v2.26 [90] and DIAMOND v2.1.8 [91], respectively. Both cDNA (parameter: –model est2genome) and protein (parameter: –model protein2genome) alignments were polished with Exonerate v2.4.0 [92], producing high quality alignments with precise intron/exon positions.

**Figure 2.**
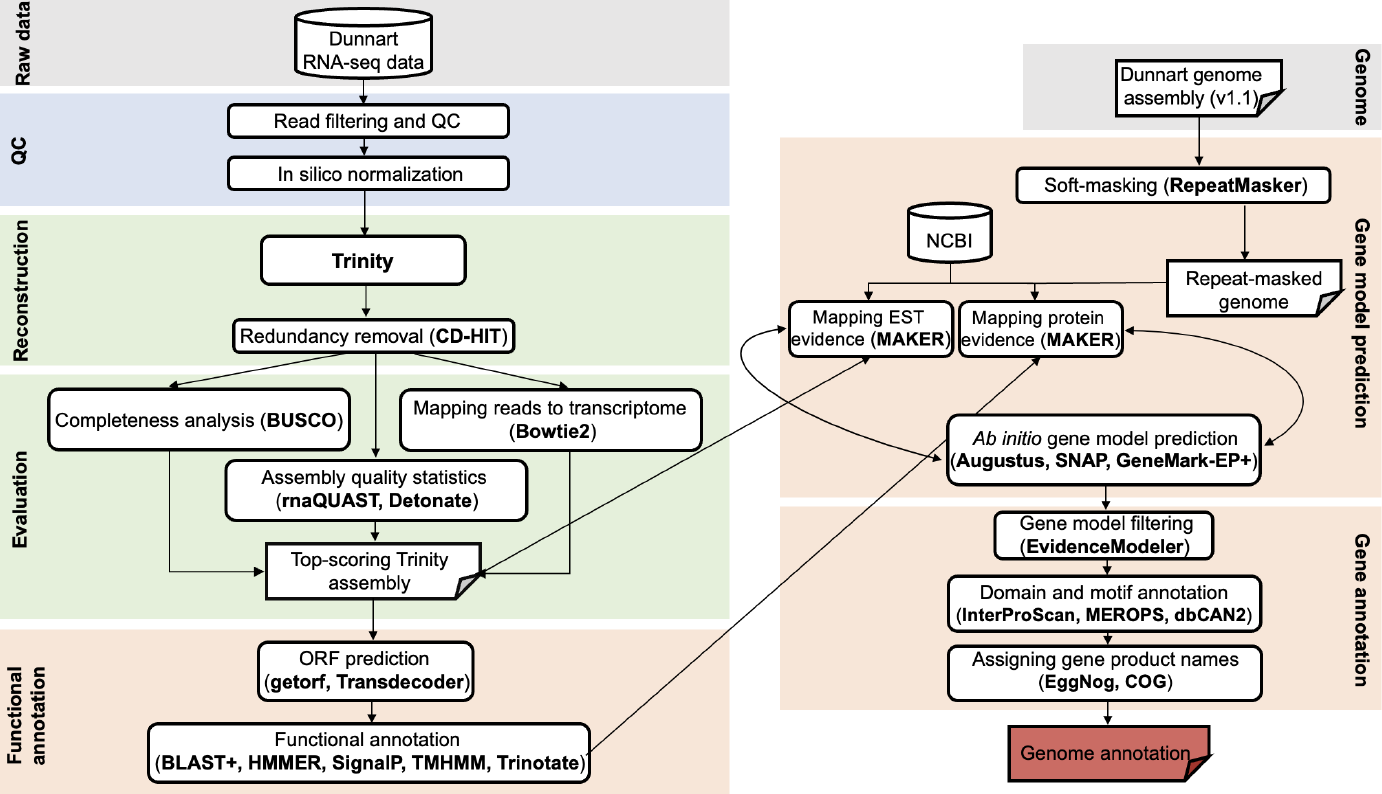
Schematic illustrating the *de novo* transcriptome generation and genome annotation workflow for the fat-tailed dunnart.

These preliminary gene models were then used to train the *ab initio* gene predictors SNAP [93], Augustus v3.4.0 [94], and GeneMark-EP+ v4.71 [95], all of which generated a statistical model representing the observed intron/exon structure in the genome. The gene model prediction process was iteratively run with MAKER2 (3 total rounds of prediction and re-training), thereby optimising the performance of the *ab initio* gene predictors. For each round, prediction quality was evaluated using BUSCO scores. Consensus gene models were identified using EVidenceModeler v2.0.0 [96], with input weights set to 2 for high-quality *ab initio* predictions, and to 1 for all other *ab initio* predictions and transcript/protein alignments. Gene models that lacked mRNA and protein homology support were excluded from the final annotation file. Lastly, gene names and putative protein functions were assigned using the aforementioned Trinotate output, as well as curated orthologous group and product names from InterProScan v5.60 [97], EggNOG v5.0 [98], MEROPS v12.4 [99], dbCAN3 v3.0.6 [100], and EuKaryotic Orthologous Groups (KOGs) [101].

## Results

To generate a genome-level annotation for the fat-tailed dunnart, we began by producing an improved draft genome assembly. We employed a hybrid approach, which integrated the ONT and PacBio long-read data with Illumina paired-end short reads [53]. This resulted in a 3.23 Gb genome that contains 1,848 scaffolds and has a scaffold N50 of 72.64 Mb. The GC content of this draft genome is 36.2% (Table 1). The genome assembly had a 94.2% recovery of complete mammalian BUSCOs.

**Table 1.** Assembly statistics for v1.0 and v1.1 of the fat-tailed dunnart genome.

|  | Genome assembly v1.0 [68] | Genome assembly v1.1 (this study) |
| --- | --- | --- |
| Genome scaffold total | 719 | 1848 |
| Genome contig total | 1154 | 2569 |
| Genome length | 2.84 Gb | 3.23 Gb |
| Genome scaffold N50 | 28.02 Mb | 72.64 Mb |
| Genome scaffold L50 | 23 | 15 |
| Genome contig N50 | 10.93 Mb | 11.19 Mb |
| Genome contig L50 | 78 | 81 |
| Max scaffold length | 442.91 Mb | 192.34 Mb |
| Max contig length | 44.29 Mb | 60.38 Mb |
| Number of scaffolds > 50 Kb | 321 | 230 |
| GC (%) | 36.25 | 36.18 |

A *de novo* reconstruction of the dunnart transcriptome was conducted using a set of 24 RNA-seq samples originating from the liver, testis, prostate, ovary, oviduct, uterus, eye, whole neonate, allantois, amnion, distal yolk sac, proximal yolk sac, and endometrium. To ensure that the most representative assembly was obtained, we sought to identify the optimal k-mer length for the Trinity contig assembly step, considering k values of 25, 29 and 32 (Table 2). Given that reference-free transcriptome assembly relies on grouping overlapping sequences of read fragments of a predetermined size (i.e., the k-mer), identifying the optimal fragment size might yield a more accurate assembly. To assess this fragment size effect, we computed multiple assembly quality metrics, including the BUSCO completeness score (transcriptome mode) and the Detonate RSEM-EVAL score for each Trinity run. The RSEM-EVAL score represents the sum of three main factors: likelihood estimates of the read representation within the assembly, the assembly prior, which assumes that each contig is generated independently, and the BIC (Bayesian Information Criterion) penalty [78]. When comparing two assemblies, a higher RSEM-EVAL score is indicative of a more complete transcriptome assembly. In our comparison, the Trinity run with a k-mer setting of 29 produced the top-scoring assembly, thus, all subsequent analysis was carried out using this assembly.

**Table 2.** Summary of the *de novo* assembly statistics for the Trinity k-mer optimization.

|  | Trinity-k25 | Trinity-k29 | Trinity-k32 |
| --- | --- | --- | --- |
| Total # of assembled Trinity transcripts | 2,588,090 | 2,093,982 | 1,960,023 |
| Mean transcript length (bp) | 731 | 830 | 922 |
| Transcript N50 (bp); E90N50 (bp) | 1193; 2990 | 1,489; 3,430 | 1,260; 3,154 |
| GC content (%) | 41.7 | 40.2 | 40.9 |
| Percentage of mapped RNA-seq PE reads (%) | 95 | 98 | 97 |
| Total BUSCO score (transcriptome mode) | C:92.1%,n:9226 | C:93.3%,n:9226 | C:93.0%,n:9226 |
| Detonate RSEM-EVAL score | -6,136.0 x 10<sup>7</sup> | -5,759.0 x 10<sup>7</sup> | -6,110.0 x 10<sup>7</sup> |

This transcriptome assembly was composed of 2,093,982 assembled transcripts (including splicing isoforms), with a GC content of 40.2% and a mean transcript length of 830 bp (Table 2). Transcript N50 was 1,489 bp, and considering only the top 90% most highly expressed transcripts (a more accurate proxy for transcriptome quality [102]) gave an E90N50 of 3,430 bp. Sample reads that were mapped back to the assembly had a very high overall alignment rate (98%) with a high percentage mapped as proper pairs (94%). In addition, the global transcriptome had a 93.3% recovery of complete mammalian BUSCOs (Mammalia_odb v10 [79]). These values are in line with, or higher than, those reported from all other available marsupial transcriptome data sets. Specifically, the global transcriptome assembly for the brown antechinus yielded 1,636,859 transcripts, with a mean length of 773 bp, transcript N50 of 1,367 bp, 96% alignment rate, and 84% complete BUSCOs [56]. The numbat global transcriptome contained 2,119,791 transcripts, a mean transcript length of 824 bp, a transcript N50 of 1,393 bp, and a BUSCO completeness score of 76.4% [58]. The Tasmanian devil transcriptome assembly consisted of 470,729 transcripts with an N50 of 687 bp and a 95% alignment rate of sample reads to the assembly [103].

Using a multi-pronged annotation approach of transcript and protein-level alignment, as well as *ab initio* gene prediction (Figure 3), we obtained 58,271 putative gene models for the fat-tailed dunnart draft genome (Table 3). Of these gene models, 21,622 were protein-coding (BLAST hits to UniProt/Swiss-Prot), which is in line with the reported gene numbers for the numbat (21,465) [58], the koala (27,669) [104], the Tasmanian devil (19,241) [46], the brown antechinus (25,111) [56], the tammar wallaby (15,290) [55], and the gray short-tailed opossum (21,384) [54] (Table 4). Furthermore, we predicted the putative function of the fat-tailed dunnart proteins using several curated protein databases (Table 3, Figure 3). We used InterProScan to identify conserved domains and assign Gene Ontology (GO) terms. A total of 24,366 transcripts were assigned InterProScan terms, and 13,507 unique genes were assigned GO terms. Specifically, GO annotations totaled 289,985, with a mean annotation level of 7.15 and a standard deviation of 2.7 (Figure 3). Running an HMMer search against the PFAM database yielded 16,308 domains, while dbCAN3 and MEROPS analyses resulted in 212 and 1,053 predictions, respectively. Altogether, these results highlight valuable avenues through which we can deepen our understanding of marsupial biology at the gene level.

**Table 3.** Fat-tailed dunnart gene and feature statistics.

|  | Fat-tailed dunnart |
| --- | --- |
| **General** |  |
| Protein-coding genes | 21,622 |
| Predicted gene models | 58,271 |
| **Transcript level** |  |
| mRNA | 50,091 |
| tRNA | 8,180 |
| Multiple exon transcripts | 44,109 |
| Single exon transcripts | 5,982 |
| Total exons | 246,391 |
| Average exon length | 146.1 |
| **Functional level** |  |
| InterProScan terms | 24,366 |
| EggNOG terms | 29,372 |
| PFAM domains | 16,308 |
| dbCAN3 (CAZymes) | 212 |
| MEROPS (proteases) | 1,053 |
| GO | 13,507 |

**Table 4.** Fat-tailed dunnart gene counts compared to the numbat, koala, Tasmanian devil, brown antechinus, tammar wallaby, gray short-tailed opossum, and eastern quoll.

|  | Number of putative genes | Number of protein-coding genes |
| --- | --- | --- |
| Fat-tailed dunnart (this study) | 58,271 | 21,622 |
| Numbat[58] | 77,806 | 21,465 |
| Koala[57] | 52,384 | 27,669 |
| Tasmanian devil[46] | 40,469 | 19,241 |
| Brown antechinus[56] | 55,827 | 25,111 |
| Tammar wallaby[55] | 122,304 | 15,290 |
| Gray short-tailed opossum[54] | 43,478 | 21,384 |
| Eastern quoll[59] | 29,622 | 14,293 |

**Figure 3.**
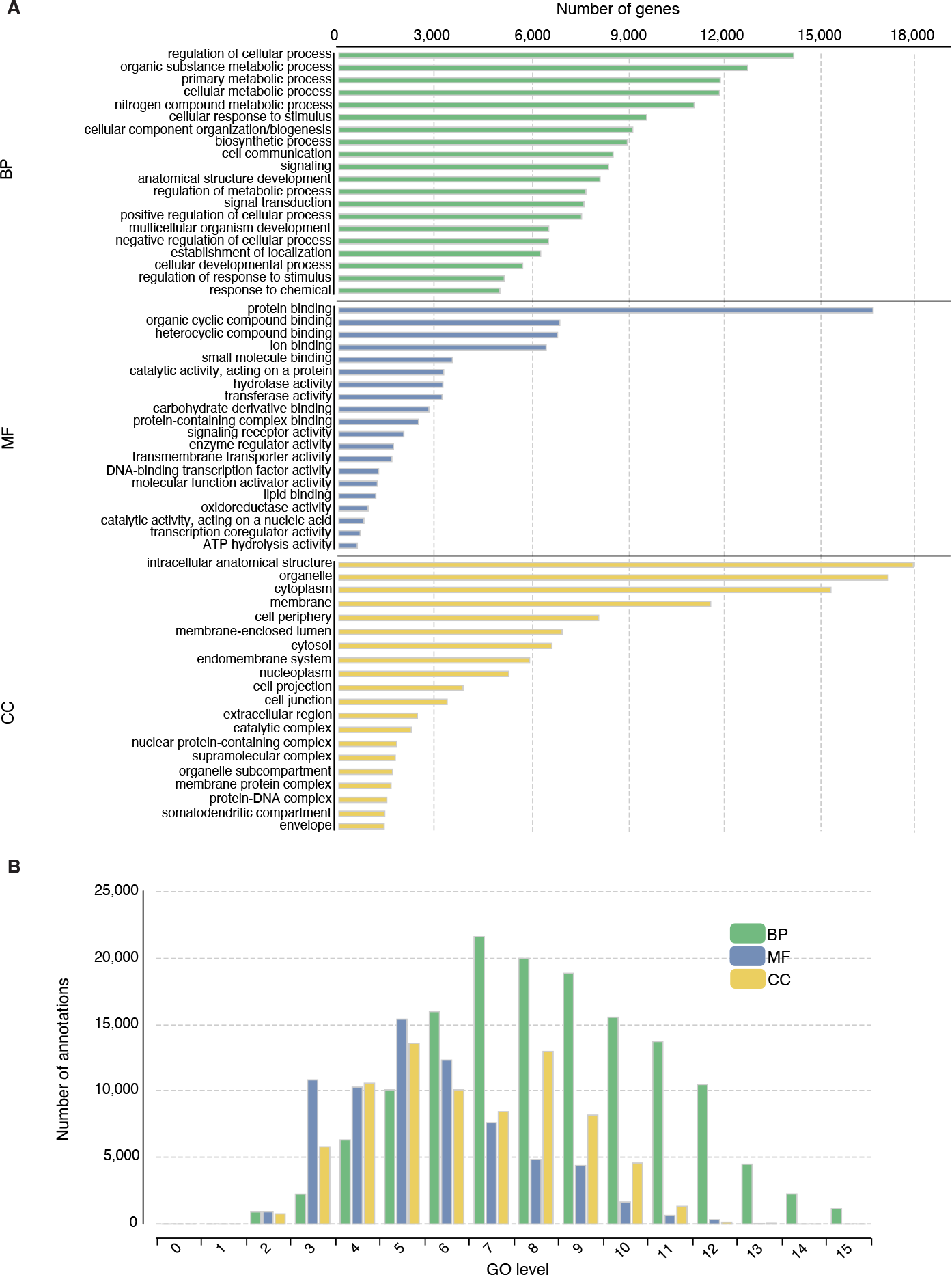
Gene ontology (GO) analysis of the fat-tailed dunnart putative genes. **(A)** GO distribution by category (at level 3) for the fat-tailed dunnart gene set. The ontology categories are BP (Biological Process), MF (Molecular Function), and CC (Cellular Component). The top 20 terms are listed for each category. **(B)** Distribution of sequence annotations for each GO level.

## Conclusion

The increased availability of genomic resources for marsupial species is critical for fostering a deeper understanding of the evolutionary history of both eutherians and marsupials. In this study, we generated a global *de novo* transcriptome assembly of the fat-tailed dunnart using RNA-seq short-read and long-read data, sampled from a diverse range of dunnart tissues. The transcriptome reconstruction contained 2,093,982 assembled transcripts, with a mean transcript length of 830 bp. The transcriptome BUSCO completeness score of 93.3% is the highest amongst all other published marsupial transcriptome BUSCOs (i.e. numbat and brown antechinus). The high overall alignment rate of reads from each of the tissues to the transcriptome (98%) further underscores that the *de novo* transcriptome is a highly accurate representation of the input reads. The dunnart draft genome annotation revealed 21,622 protein-coding genes, in line with previously reported marsupial gene counts. Overall, these resources provide novel insights into the unique genomic architecture of the fat-tailed dunnart, and will therefore serve as valuable tools for future comparative mammalian studies.

## Data and Code Availability

The fat-tailed dunnart transcriptome, draft genome, and genome annotation are available through Figshare https://melbourne.figshare.com/account/home#/projects/183307. The scripts for reproducing the genome annotation workflow have been made available here: https://gitlab.svi.edu.au/igr-lab/dunnart_genome_annotation. All raw sequencing reads have been deposited at the National Center for Biotechnology Information (NCBI) Sequence Read Archive under the accession number PRJNA1028148.

## Declarations

## List of abbreviations

BLAST: Basic Local Alignment Search Tool
bp: base pair
BUSCO: Benchmarking Universal Single-Copy Orthologs
CDS: coding sequences
Gb: Gigabase
Kb: Kilobase
Mb: Megabase
NCBI: National Center for Biotechnology Information
ONT: Oxford Nanopore Technologies
PacBio: Pacific Biosciences
PE: paired-end
RNA-seq: RNA sequencing

## Ethics Statement

All sample collection was approved by the University of Melbourne Animal Ethics Committee (Project ID 10206).

## Author’s Contributions

SF, AJP, CYF, and IGR conceived the project. SF collected and prepared the samples. NI assembled and annotated the global transcriptome. CYF assembled the draft genome. NI annotated the draft genome. NI drafted the manuscript with edits from all authors. All authors read and approved the final version of the manuscript.

## Acknowledgements

NI was supported by the University of Melbourne Research Training Program Scholarship, the Rowden White Scholarship, the Dame Margaret Blackwood Soroptimist Scholarship, and St. Vincent’s Institute Top-up Scholarship. St Vincent’s Institute acknowledges the infrastructure support it receives from the National Health and Medical Research Council Independent Research Institutes Infrastructure Support Program and from the Victorian Government through its Operational Infrastructure Support Program.

## References

1. Commons W, Commons W, editor, Fat-tailed Dunnart area. Wikimedia Commons; 2010. https://commons.wikimedia.org/wiki/File:Fat-tailed_Dunnart_area.png, iUCN Red List of Threatened Species, CC BY-SA 3.0.

2. Duchêne DA, Bragg JG, Duchêne S, Neaves LE, Potter S, Moritz C, et al. Analysis of Phylogenomic Tree Space Resolves Relationships Among Marsupial Families. Syst Biol 2018 May;67(3):400–412.

3. Doronina L, Feigin CY, Schmitz J. Reunion of Australasian Possums by Shared SINE Insertions. Syst Biol 2022 Aug;71(5):1045–1053.

4. Jackson SM, Jackson S, Groves C. Taxonomy of Australian Mammals. Csiro Publishing; 2015.

5. Wilson DE, Reeder DM. Mammal Species of the World: A Taxonomic and Geographic Reference. JHU Press; 2005.

6. Archer M, Beck R, Gott M, Hand S, Godthelp H, Black K. Australia’s first fossil marsupial mole (Notoryctemorphia) resolves controversies about their evolution and palaeoenvironmental origins. Proceedings of the Royal Society B: Biological Sciences 2010 Nov;278(1711):1498–1506.

7. Diogo R, Bello-Hellegouarch G, Kohlsdorf T, Esteve-Altava B, Molnar JL. Comparative myology and evolution of marsupials and other vertebrates, with notes on complexity, Bauplan, and “scala naturae”. Anat Rec 2016 Sep;299(9):1224–1255.

8. Stein BR. Comparative limb myology of two opossums, Didelphis and Chironectes. J Morphol 1981 Jul;169(1):113–140.

9. Schmitz J, Ohme M, Suryobroto B, Zischler H. The colugo (Cynocephalus variegatus, Dermoptera): the primates’ gliding sister? Mol Biol Evol 2002 Dec;19(12):2308–2312.

10. Casanovas-Vilar I, Garcia-Porta J, Fortuny J, Sanisidro ó, Prieto J, Querejeta M, et al. Oldest skeleton of a fossil flying squirrel casts new light on the phylogeny of the group. Elife 2018 Oct;7.

11. Henneberg M, Lambert K, Leigh C. Fingerprint homoplasy: koalas and humans. Natural Science 1997;.

12. McGhee GR. Convergent Evolution: Limited Forms Most Beautiful. MIT Press; 2011.

13. Freeman C. 4. A Tasmanian Wolf. In: Paper Tiger Brill; 2010. p. 117–149.

14. Feigin CY, Newton AH, Pask AJ. Widespread cis-regulatory convergence between the extinct Tasmanian tiger and gray wolf. Genome Res 2019 Oct;29(10):1648–1658.

15. Geiser F, Körtner G, Schmidt I. Leptin increases energy expenditure of a marsupial by inhibition of daily torpor. American Journal of Physiology-Regulatory, Integrative and Comparative Physiology 1998 Nov;275(5):R1627–R1632.

16. Hing S, Narayan E, Thompson RCA, Godfrey S. A review of factors influencing the stress response in Australian marsupials. Conserv Physiol 2014 Jul;2(1):cou027.

17. Karlen SJ, Krubitzer L. The functional and anatomical organization of marsupial neocortex: evidence for parallel evolution across mammals. Prog Neurobiol 2007 Jun;82(3):122–141.

18. Krause WJ, Cutts JH, Leeson CR. Postnatal development of the epidermis in a marsupial, Didelphis virginiana. J Anat 1978;.

19. Cutts JH, Leeson CR, Krause WJ. The postnatal development of the liver in a marsupial, Didelphis virginiana. 1. Light microscopy. J Anat 1973 Sep;115(Pt 3):327–346.

20. Fadem BH, Harder JD. Evidence for High Levels of Androgen in Peripheral Plasma during Postnatal Development in a Marsupial: The Gray Short-tailed Opossum (Monodeiphis Domestica)1. Biol Reprod 1992 Jan;46(1):105–108.

21. Runciman SI, Baudinette RV, Gannon BJ. Postnatal development of the lung parenchyma in a marsupial: the tammar wallaby. Anat Rec 1996 Feb;244(2):193–206.

22. Goldingay RL. The behavioural ecology of the gliding marsupial, Petaurus australis. PhD thesis, University of Wollongong; 1989.

23. Menário Costa W, King WJ, Bonnet T, Festa-Bianchet M, Kruuk LEB. Early-life behavior, survival, and maternal personality in a wild marsupial. Behav Ecol 2023 Sep;p. arad070.

24. Russell EM. Social behaviour and social organization of marsupials. Mamm Rev 1984 Sep;14(3):101–154.

25. Renfree MB. Monotreme and marsupial reproduction. Reprod Fertil Dev 1995;7(5):1003–1020.

26. Sharman GB. Reproductive physiology of marsupials. Science 1970 Feb;167(3922):1221–1228.

27. Harder JD, Jackson LM. Chemical communication and reproduction in the gray short-tailed opossum (Monodelphis domestica). Vitam Horm 2010;83:373–399.

28. Bergallo HG, Cerqueira R. Reproduction and growth of the opossumMonodelphis domestica(Mammalia: Didelphidae) in northeastern Brazil. J Zool 1994 Apr;232(4):551–563.

29. Chen Y, Yu H, Pask AJ, Fujiyama A, Suzuki Y, Sugano S, et al. Hormone-responsive genes in the SHH and WNT/β-catenin signaling pathways influence urethral closure and phallus growth. Biol Reprod 2018 Oct;99(4):806–816.

30. Coveney D, Shaw G, Hutson JM, Renfree MB. Effect of an anti-androgen on testicular descent and inguinal closure in a marsupial, the tammar wallaby (Macropus eugenii). Reproduction 2002 Dec;124(6):865–874.

31. Moore HDM, Thurstan SM. Sexual differentiation in the grey short-tailed opossum, Monodelphis domestica, and the effect of oestradiol benzoate on development in the male. J Zool 1990 Aug;221(4):639–658.

32. Renfree MB, Pask AJ, Shaw G. Sex down under: the differentiation of sexual dimorphisms during marsupial development. Reprod Fertil Dev 2001;13(7-8):679–690.

33. Pask AJ, Harry JL, Renfree MB, Marshall Graves JA. Absence of SOX3 in the developing marsupial gonad is not consistent with a conserved role in mammalian sex determination. Genesis 2000 Aug;27(4):145–152.

34. Pask A, Renfree MB, Marshall Graves JA. The human sex-reversing ATRX gene has a homologue on the marsupial Y chromosome, ATRY: implications for the evolution of mammalian sex determination. Proc Natl Acad Sci U S A 2000 Nov;97(24):13198–13202.

35. Scherer G, Schmid M. Genes and mechanisms in vertebrate sex determination. Introduction. EXS 2001;91(3):XI–XII.

36. Hornecker JL, Samollow PB, Robinson ES, Vandeberg JL, McCarrey JR. Meiotic sex chromosome inactivation in the marsupial Monodelphis domestica. Genesis 2007 Nov;45(11):696–708.

37. Foster JW, Brennan FE, Hampikian GK, Goodfellow PN, Sinclair AH, Lovell-Badge R, et al. Evolution of sex determination and the Y chromosome: SRY-related sequences in marsupials. Nature 1992 Oct;359(6395):531–533.

38. Ishihara T, Hickford D, Shaw G, Pask AJ, Renfree MB. DNA methylation dynamics in the germline of the marsupial tammar wallaby, Macropus eugenii. DNA Res 2019 Feb;26(1):85–94.

39. Whitworth DJ, Pask AJ. The X factor: X chromosome dosage compensation in the evolutionarily divergent monotremes and marsupials. Semin Cell Dev Biol 2016 Aug;56:117–121.

40. Wang X, Douglas KC, Vandeberg JL, Clark AG, Samollow PB. Chromosome-wide profiling of X-chromosome inactivation and epigenetic states in fetal brain and placenta of the opossum, Monodelphis domestica. Genome Res 2014 Jan;24(1):70–83.

41. Das R, Anderson N, Koran MI, Weidman JR, Mikkelsen TS, Kamal M, et al. Convergent and divergent evolution of genomic imprinting in the marsupial Monodelphis domestica. BMC Genomics 2012 Aug;13:394.

42. McAllan BM. Dasyurid marsupials as models for the physiology of ageing in humans. Aust J Zool 2006 Jun;54(3):159–172.

43. Bartkowska K, Tepper B, Turlejski K, Djavadian R. Postnatal and Adult Neurogenesis in Mammals, Including Marsupials. Cells 2022 Sep;11(17).

44. Szdzuy K, Zeller U, Renfree M, Tzschentke B, Janke O. Postnatal lung and metabolic development in two marsupial and four eutherian species. J Anat 2008 Feb;212(2):164–179.

45. Cook LE, Newton AH, Hipsley CA, Pask AJ. Postnatal development in a marsupial model, the fat-tailed dunnart (Sminthopsis crassicaudata; Dasyuromorphia: Dasyuridae). Commun Biol 2021 Sep;4(1):1028.

46. Stammnitz MR, Gori K, Kwon YM, Harry E, Martin FJ, Billis K, et al. The evolution of two transmissible cancers in Tasmanian devils. Science 2023 Apr;380(6642):283–293.

47. De Leo AA. Genome Evolution in Australian Marsupials. University of Melbourne, Department of Zoology, Faculty of Science; 2005.

48. Graves JA. Mammalian genome evolution: new clues from comparisons of eutherians, marsupials and monotremes. Comp Biochem Physiol A Comp Physiol 1991;99(1-2):5–11.

49. Deakin JE, O’Neill RJ. Evolution of Marsupial Genomes. Annu Rev Anim Biosci 2020 Feb;8:25–45.

50. Ishihara T, Hickford D, Fenelon JC, Griffith OW, Suzuki S, Renfree MB. Evolution of the Short Form of DNMT3A, DNMT3A2, Occurred in the Common Ancestor of Mammals. Genome Biol Evol 2022 Jul;14(7).

51. Deakin JE. Marsupial genome sequences: providing insight into evolution and disease. Scientifica 2012 Nov;2012:543176.

52. Janke A, Feldmaier-Fuchs G, Thomas WK, von Haeseler A, Pääbo S. The marsupial mitochondrial genome and the evolution of placental mammals. Genetics 1994 May;137(1):243–256.

53. Feigin C, Frankenberg S, Pask A. A Chromosome-Scale Hybrid Genome Assembly of the Extinct Tasmanian Tiger (Thylacinus cynocephalus). Genome Biol Evol 2022 Apr;14(4).

54. Mikkelsen TS, Wakefield MJ, Aken B, Amemiya CT, Chang JL, Duke S, et al. Genome of the marsupial Monodelphis domestica reveals innovation in non-coding sequences. Nature 2007 May;447(7141):167–177.

55. Renfree MB, Papenfuss AT, Deakin JE, Lindsay J, Heider T, Belov K, et al. Genome sequence of an Australian kangaroo, Macropus eugenii, provides insight into the evolution of mammalian reproduction and development. Genome Biol 2011 Aug;12(8):R81.

56. Brandies PA, Tang S, Johnson RSP, Hogg CJ, Belov K. The first Antechinus reference genome provides a resource for investigating the genetic basis of semelparity and age-related neuropathologies. GigaByte 2020 Nov;2020:gigabyte7.

57. Johnson RN, O’Meally D, Chen Z, Etherington GJ, Ho SYW, Nash WJ, et al. Adaptation and conservation insights from the koala genome. Nat Genet 2018;50(8):1102–1111.

58. Peel E, Silver L, Brandies P, Hayakawa T, Belov K, Hogg CJ. Genome assembly of the numbat (Myrmecobius fasciatus), the only termitivorous marsupial. GigaByte 2022 Mar;2022:gigabyte47.

59. Hartley GA, Frankenberg SR, Robinson NM, MacDonald AJ, Hamede RK, Burridge CP, et al. Reference genome of the endangered eastern quoll (Dasyurus viverrinus); 2023, unpublished.

60. Polymeropoulos ET, Jastroch M, Frappell PB. Absence of adaptive nonshivering thermogenesis in a marsupial, the fat-tailed dunnart (Sminthopsis crassicaudata). J Comp Physiol B 2012 Apr;182(3):393–401.

61. Suárez R, Paolino A, Kozulin P, Fenlon LR, Morcom LR, Englebright R, et al. Development of body, head and brain features in the Australian fat-tailed dunnart (Sminthopsis crassicaudata; Marsupialia: Dasyuridae); A postnatal model of forebrain formation. PLoS One 2017 Sep;12(9):e0184450.

62. Garrett A, Lannigan V, Yates NJ, Rodger J, Mulders W. Physiological and anatomical investigation of the auditory brainstem in the Fat-tailed dunnart (Sminthopsis crassicaudata). PeerJ 2019 Sep;7:e7773.

63. Noy EB, Scott MK, Grommen SVH, Robert KA, De Groef B. Molecular cloning and tissue distribution of Crh and Pomc mRNA in the fat-tailed dunnart (Sminthopsis crassicaudata), an Australian marsupial. Gene 2017 Sep;627:26–31.

64. Suárez R, Paolino A, Kozulin P, Fenlon LR, Morcom LR, Englebright R, et al. Development of body, head and brain features in the Australian fat-tailed dunnart (Sminthopsis crassicaudata; Marsupialia: Dasyuridae); A postnatal model of forebrain formation. PLoS One 2017 Sep;12(9):e0184450.

65. Collins LR. Monotremes and marsupials. Smithsonian Institution 1973;.

66. Tyndale-Biscoe CH, Janssens PA. The Developing Marsupial: Models for Biomedical Research. Springer Science & Business Media; 2012.

67. Godfrey GK, Crowcroft P. Breeding the Fat-tailed marsupial mouse in captivity. Int Zoo Yearbook 1971 Jan;11(1):33–38.

68. Cook LE, Feigin CY, Pask AJ, Romero IG. Cis-regulatory landscapes of the fat-tailed dunnart and mouse provide insights into the drivers of craniofacial heterochrony; 2023, unpublished.

69. Kolmogorov M, Yuan J, Lin Y, Pevzner PA. Assembly of long, error-prone reads using repeat graphs. Nat Biotechnol 2019 May;37(5):540–546.

70. Guan D, McCarthy SA, Wood J, Howe K, Wang Y, Durbin R. Identifying and removing haplotypic duplication in primary genome assemblies. Bioinformatics 2020 May;36(9):2896–2898.

71. Coombe L, Li JX, Lo T, Wong J, Nikolic V, Warren RL, et al. LongStitch: high-quality genome assembly correction and scaffolding using long reads. BMC Bioinformatics 2021 Oct;22(1):534.

72. Walker BJ, Abeel T, Shea T, Priest M, Abouelliel A, Sakthikumar S, et al. Pilon: an integrated tool for comprehensive microbial variant detection and genome assembly improvement. PLoS One 2014 Nov;9(11):e112963.

73. Bolger AM, Lohse M, Usadel B. Trimmomatic: a flexible trimmer for Illumina sequence data. Bioinformatics 2014 Aug;30(15):2114–2120.

74. Vasimuddin M, Misra S, Li H, Aluru S. Efficient Architecture-Aware Acceleration of BWA-MEM for Multicore Systems. In: 2019 IEEE International Parallel and Distributed Processing Symposium (IPDPS); 2019. p. 314–324.

75. Li H, Handsaker B, Wysoker A, Fennell T, Ruan J, Homer N, et al. The Sequence Alignment/Map format and SAMtools. Bioinformatics 2009 Aug;25(16):2078–2079.

76. Andrews S. FastQC: a quality control tool for high throughput sequence data. Babraham Bioinformatics 2010;.

77. Haas BJ, Papanicolaou A, Yassour M, Grabherr M, Blood PD, Bowden J, et al. De novo transcript sequence reconstruction from RNA-seq using the Trinity platform for reference generation and analysis. Nat Protoc 2013 Aug;8(8):1494–1512.

78. Li B, Fillmore N, Bai Y, Collins M, Thomson JA, Stewart R, et al. Evaluation of de novo transcriptome assemblies from RNA-Seq data. Genome Biol 2014 Dec;15(12):553.

79. Simão FA, Waterhouse RM, Ioannidis P, Kriventseva EV, Zdobnov EM. BUSCO: assessing genome assembly and annotation completeness with single-copy orthologs. Bioinformatics 2015 Oct;31(19):3210–3212.

80. Langmead B, Salzberg SL. Fast gapped-read alignment with Bowtie 2. Nat Methods 2012 Mar;9(4):357–359.

81. Huang Y, Niu B, Gao Y, Fu L, Li W. CD-HIT Suite: a web server for clustering and comparing biological sequences. Bioinformatics 2010 Mar;26(5):680–682.

82. Camacho C, Coulouris G, Avagyan V, Ma N, Papadopoulos J, Bealer K, et al. BLAST+: architecture and applications. BMC Bioinformatics 2009 Dec;10:421.

83. Boutet E, Lieberherr D, Tognolli M, Schneider M, Bairoch A. UniProtKB/Swiss-Prot. Methods Mol Biol 2007;406:89–112.

84. Finn RD, Clements J, Eddy SR. HMMER web server: interactive sequence similarity searching. Nucleic Acids Res 2011 Jul;39(Web Server issue):W29–37.

85. Mistry J, Chuguransky S, Williams L, Qureshi M, Salazar GA, Sonnhammer ELL, et al. Pfam: The protein families database in 2021. Nucleic Acids Res 2021 Jan;49(D1):D412–D419.

86. Teufel F, Almagro Armenteros JJ, Johansen AR, Gíslason MH, Pihl SI, Tsirigos KD, et al. SignalP 6.0 predicts all five types of signal peptides using protein language models. Nat Biotechnol 2022 Jul;40(7):1023–1025.

87. Hallgren J, Tsirigos KD, Pedersen MD, Armenteros JJA, Marcatili P, Nielsen H, et al. DeepTMHMM predicts alpha and beta transmembrane proteins using deep neural networks; 2022, unpublished.

88. Nishimura D. RepeatMasker. Biotech Software & Internet Report 2000 Apr;1(1-2):36–39.

89. Holt C, Yandell M. MAKER2: an annotation pipeline and genome-database management tool for second-generation genome projects. BMC Bioinformatics 2011 Dec;12:491.

90. Li H. Minimap2: pairwise alignment for nucleotide sequences. Bioinformatics 2018 Sep;34(18):3094–3100.

91. Buchfink B, Xie C, Huson DH. Fast and sensitive protein alignment using DIAMOND. Nat Methods 2015 Jan;12(1):59–60.

92. Slater GSC, Birney E. Automated generation of heuristics for biological sequence comparison. BMC Bioinformatics 2005 Feb;6:31.

93. Korf I. Gene finding in novel genomes. BMC Bioinformatics 2004 May;5:59.

94. Stanke M, Diekhans M, Baertsch R, Haussler D. Using native and syntenically mapped cDNA alignments to improve de novo gene finding. Bioinformatics 2008 Mar;24(5):637–644.

95. Brůna T, Lomsadze A, Borodovsky M. GeneMark-EP+: eukaryotic gene prediction with self-training in the space of genes and proteins. NAR Genom Bioinform 2020 Jun;2(2):lqaa026.

96. Haas BJ, Salzberg SL, Zhu W, Pertea M, Allen JE, Orvis J, et al. Automated eukaryotic gene structure annotation using EVidenceModeler and the Program to Assemble Spliced Alignments. Genome Biol 2008 Jan;9(1):R7.

97. Jones P, Binns D, Chang HY, Fraser M, Li W, McAnulla C, et al. InterProScan 5: genome-scale protein function classification. Bioinformatics 2014 May;30(9):1236–1240.

98. Huerta-Cepas J, Szklarczyk D, Heller D, Hernández-Plaza A, Forslund SK, Cook H, et al. eggNOG 5.0: a hierarchical, functionally and phylogenetically annotated orthology resource based on 5090 organisms and 2502 viruses. Nucleic Acids Res 2019 Jan;47(D1):D309–D314.

99. Rawlings ND, Barrett AJ, Bateman A. MEROPS: the peptidase database. Nucleic Acids Res 2010 Jan;38(Database issue):D227–33.

100. Zheng J, Ge Q, Yan Y, Zhang X, Huang L, Yin Y. dbCAN3: automated carbohydrate-active enzyme and substrate annotation. Nucleic Acids Res 2023 Jul;51(W1):W115–W121.

101. Tatusov RL, Fedorova ND, Jackson JD, Jacobs AR, Kiryutin B, Koonin EV, et al. The COG database: an updated version includes eukaryotes. BMC Bioinformatics 2003 Sep;4:41.

102. Haas B. Transcriptome contig Nx and ExN50 stats. Transcriptome Contig Nx and ExN50 stats 2016;.

103. Murchison EP, Schulz-Trieglaff OB, Ning Z, Alexandrov LB, Bauer MJ, Fu B, et al. Genome sequencing and analysis of the Tasmanian devil and its transmissible cancer. Cell 2012 Feb;148(4):780–791.

104. Blanchard AM, Emes RD, Greenwood AD, Holmes N, Loose MW, McEwen GK, et al. Genome Reference Assembly for Bottlenecked Southern Australian Koalas. Genome Biol Evol 2023 Jan;15(1).

